# Physiological and Metabolomic Consequences of Reduced Expression of the Drosophila *brummer* Triglyceride Lipase

**DOI:** 10.1101/2021.08.04.455099

**Authors:** Nestor O. Nazario-Yepiz, Jaime Fernández Sobaberas, Roberta Lyman, Marion R. Campbell, Vijay Shankar, Robert R. H. Anholt, Trudy F. C. Mackay

## Abstract

Disruption of lipolysis has widespread effects on intermediary metabolism and organismal phenotypes. Defects in lipolysis can be modeled in *Drosophila melanogaster* through genetic manipulations of *brummer* (*bmm*), which encodes a triglyceride lipase orthologous to mammalian Adipose Triglyceride Lipase. RNAi-mediated knock-down of *bmm* in all tissues or metabolic specific tissues results in reduced locomotor activity, altered sleep patterns and reduced lifespan. Metabolomic analysis on flies in which *bmm* is downregulated reveals a marked reduction in medium chain fatty acids, long chain saturated fatty acids and long chain monounsaturated and polyunsaturated fatty acids, and an increase in diacylglycerol levels. Elevated carbohydrate metabolites and tricarboxylic acid intermediates indicate that impairment of fatty acid mobilization as an energy source may result in upregulation of compensatory carbohydrate catabolism. *bmm* downregulation also results in elevated levels of serotonin and dopamine neurotransmitters, possibly accounting for the impairment of locomotor activity and sleep patterns. Physiological phenotypes and metabolomic changes upon reduction of *bmm* expression show extensive sexual dimorphism. Altered metabolic states in the Drosophila model are relevant for understanding human metabolic disorders, since pathways of intermediary metabolism are conserved across phyla.

## Introduction

Metabolic syndrome and metabolic diseases impact a large proportion of the world population: a global analysis of 195 countries showed that 604 million adults and 108 million children had obesity in 2015 [1, 2]. Although diets with poor ratios of nutrients coupled with a reduction of physical activity contribute to metabolic diseases, genetic risk alleles in different populations are also major contributors [3]. Obesity can impair cognition and increase the risk for psychiatric conditions [4], but the link between obesity and brain function is unclear [5].

*Drosophila* provides a powerful model system for comprehensive analyses of physiological, behavioral and metabolic consequences of disruption of intermediary metabolism through selective tissue-specific disruption of target genes under controlled dietary conditions [6–9]. Comprehensive studies in *Drosophila* are further facilitated through the public availability of a wide array of genetic resources that can facilitate in-depth systems genetic studies of metabolic regulation [10–15]. *Drosophila* has organ systems analogous to those of mammals that control the uptake, storage and metabolism of nutrients [16]. Digestion and absorption of lipids occur in the midgut, from where lipids are transported by lipoproteins through the hemolymph to other organs [17]. Following feeding, the fat body stores lipids taken up from the hemolymph as triacylglycerides (TAGs) and cholesterol esters. This organ performs lipogenesis by converting carbohydrates via the glycolytic pathway into TAGs and lipolysis to release fatty acids from TAGs when energy is needed. Oenocytes, specialized hepatocyte-homologous cells in Drosophila, are required with the fat body for regulation of lipid mobilization [18].

Lipids in the fat body are accumulated in lipid droplets. Their release is controlled by adipokinetic hormone (Akh), the functional counterpart of glucagon in humans, via a G protein-coupled Akh receptor (Akh-R) pathway [19, 20]. Akh-R signaling uses a canonical cAMP/PKA signal transduction pathway to regulate the phosphorylation of Perlipin1, which allows access to TAG lipases in lipid droplets [21]. Activation of Akh-R signaling also regulates the expression of the triglyceride lipase *brummer* (*bmm*) [9, 22].

*bmm* is the ortholog of the human *PNPLA2* gene that encodes Adipose Triglyceride Lipase (ATGL), the major mammalian TAG lipase. ATGL hydrolyzes TAGs at the sn-1 and sn-2 positions to release fatty acids for catabolism. Homozygous *bmm* null alleles exhibit embryonic lethality, but some “escapers” reach the adult stage and present excessive fat storage [23]. Humans with mutations in the *bmm* ortholog *PNPLA2* have neutral lipid storage disease with myopathy (NLSDM), characterized by abnormal accumulation of fat in different tissues [24]. Regulation of the balance between lipid storage and breakdown through *bmm* is sexually dimorphic and under neural control [25]. Defects in lipid metabolism are associated with various neurodegenerative diseases in *Drosophila* [26], but little is known about the behavioral phenotypes and physiological effects of *bmm* downregulation.

Here, we show that RNAi-mediated suppression of *bmm* results in impaired locomotion and altered sleep patterns. Metabolomic analysis shows that these effects are accompanied by shifts in intermediary metabolism and changes in levels of neurotransmitters in the brain.

## Results

### RNAi-mediated reduction in *bmm* expression

We used targeted RNA interference to reduce the expression of *bmm*. We employed in all experiments only one copy of the *Gal4* driver and *UAS-bmm-RNAi* to generate hypomorphic effects. First, we analyzed *bmm* transcript levels with ubiquitous expression of *bmm-RNAi* in adult flies using RT-qPCR. We used two different *UAS-bmm-RNAi* constructs, *bmm-RNAi^V37877^* and *bmm-RNAi^V37880^*. For both *Ubi > bmm-RNAi* flies, levels of *bmm* transcripts were significantly lower compared to their controls for both sexes, but the effect of *Ubi > bmm-RNAi^V37877^* was more pronounced than the effect of *Ubi > bmm-RNAi^V37880^*. The reduction of *bmm* transcripts was greater in males than in females for both genotypes (Fig. 1a and Table S1). Both RNAi constructs affected the same phenotypes, described below, but the effects were quantitatively different due to differences in the magnitude of *bmm* knockdown. For example, *Ubi > bmm-RNAi^V37877^* males, which showed the greatest reduction in *bmm* transcript, exhibited a phenotype of spread wings, which was not observed in other knockdown genotypes (Fig. 1b).

**Fig 1.**
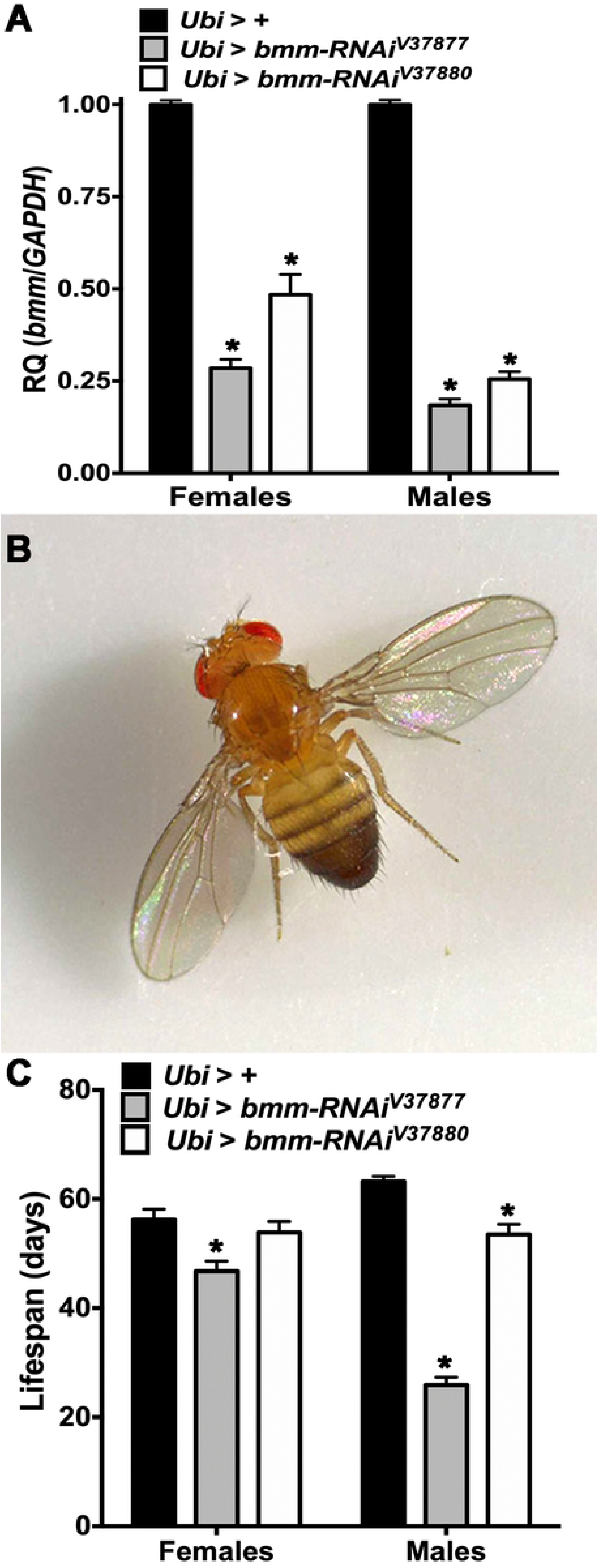
RT-qPCR and lifespan of *Ubi* > *bmm-RNAi* flies. Offspring of *Ubi-Gal4* mated with GD control (*Ubi* > +) or *UAS-bmm-RNAi* flies (*Ubi* > *bmm-RNAi^V37877^* and *Ubi* > *bmm-RNAi^V37880^*) were used for the assays. Asterisks indicate significant differences at *p* < 0.05 compared with the appropriate female or male control, following Tukey’s correction for multiple tests. (a) RT-qPCR (n=9 for females and males of each genotype), relative quantification (RQ) was done using the 2**^-ΔΔCt^** method, and *GAPDH* was used as an internal standard. (b) Spread wings phenotype in males of *Ubi* > *bmm-RNAi^V37877^* (100%, n=100). (c) Average lifespan for *Ubi* > + (n=132 for females and 118 for males), *Ubi* > *bmm-RNAi^V37877^* (n=136 for females and 134 for males) and *Ubi* > *bmm-RNAi^V37880^* (n=137 for females and 132 for males). Error bars are standard errors of the mean (SEM). ANOVA tests are reported in Table S1.

### *Ubi* > *bmm-RNAi* flies have reduced lifespan

To corroborate previously reported effects on lifespan of *bmm* mutant escapers [23], we measured lifespan of *Ubi > bmm-RNAi* flies under standard culture conditions. Females of *Ubi > bmm-RNAi^V37877^* and males of both *Ubi > bmm-RNAi* lines had reduced lifespan in a sex biased manner (Fig. 1c and Table S1). Moreover, penetrance of this phenotype matched perfectly with the titration of *bmm* transcript level of *Ubi > bmm-RNAi* flies. The level of *bmm* transcript of our allelic series ranged from high hypomorphic to low hypomorphic as follows: *Ubi > bmm-RNAi^V37877^* males, *Ubi > bmm-RNAi^V37880^* males, *Ubi > bmm-RNAi^V37877^* females and *Ubi > bmm-RNAi^V37880^* females.

### *bmm* downregulation reduces locomotor activity

We used locomotor activity as a phenotypic read-out for impaired energy metabolism as a consequence of reduced triglyceride access due to *bmm* down regulation. We recorded continuous activity of *Ubi > bmm-RNAi* flies for one week using the Drosophila Activity Monitor (DAM) system. The 24h locomotor activity was significantly lower in *Ubi > bmm-RNAi^V37877^* males than control males (Fig. 2a). Locomotor activity was reduced both during the daytime and the nighttime (Fig. 2b), which is apparent from their average activity profiles (see also Fig. 2c and Table S2).

**Fig 2.**
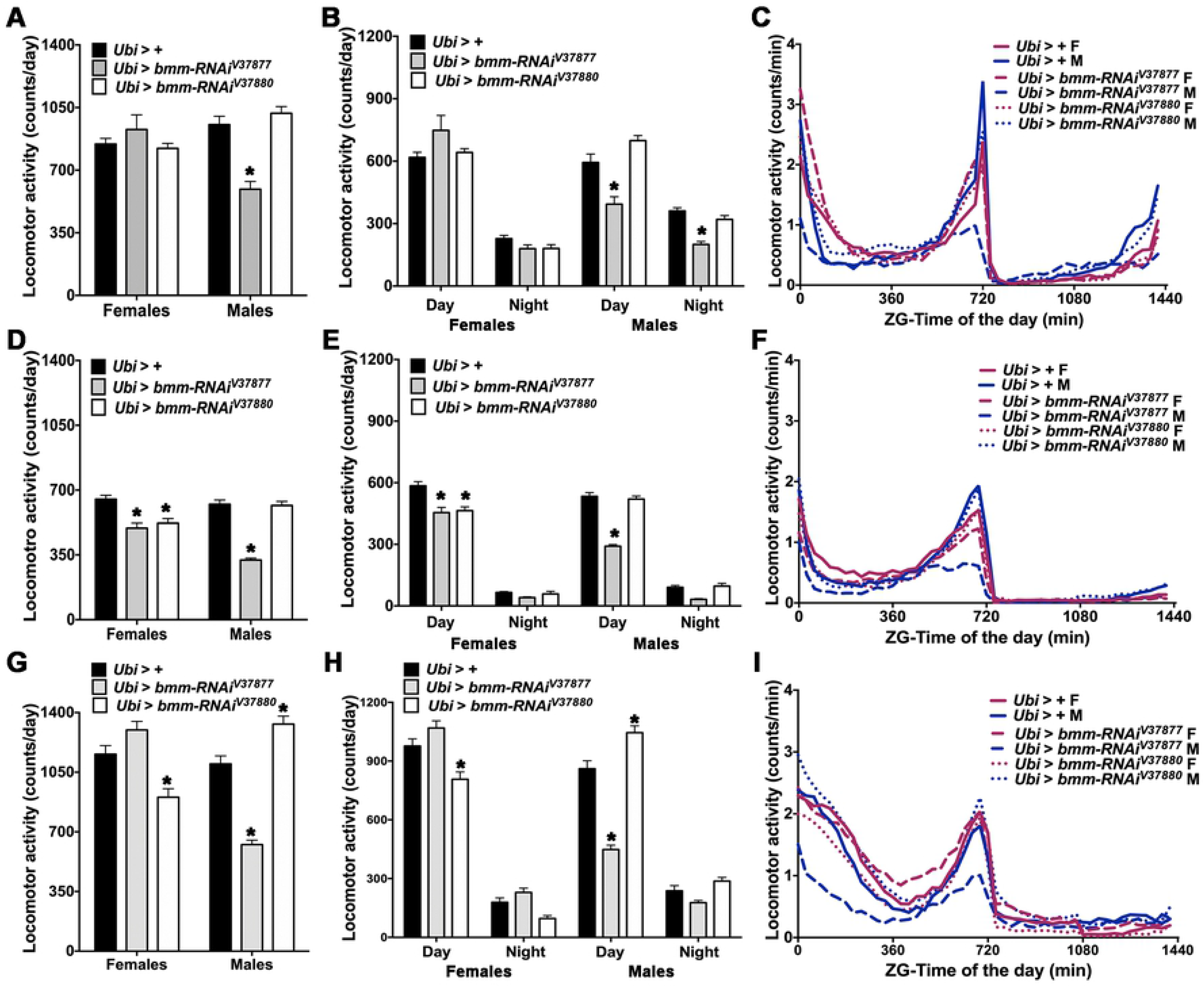
Locomotor activity in *Ubi* > *bmm-RNAi* flies. Offspring of *Ubi-Gal4* mated with GD control (*Ubi* > +) or *UAS-bmm-RNAi* flies (*Ubi* > *bmm-RNAi^V37877^* and *Ubi* > *bmm-RNAi^V37880^*) were used for the locomotor activity assay in normal feeding (a-c), normal feeding without wings (d-f) and starvation conditions (g-i). (a, d and g) Average of daily locomotor activity. (b, e and h) Average of locomotor activity during the daytime and nighttime. (c, f and i) Average activity profiles in Zeitgeber (ZG) time. Averages were calculated from 7 days of the behavior assays except for starvation, which was measured over 3 days. For this figure and Fig. 3, “n” for normal feeding in females (F) and males (M) were: *Ubi* > + F (n=49), *Ubi* > + M (n=57), *Ubi* > *bmm-RNAi^V37877^* F (n=54), *Ubi* > *bmm-RNAi^V37877^* M (n=53), *Ubi > bmm-RNAi^V37880^* F (n=57) and *Ubi > bmm-RNAi^V37880^* M (n=62); “n” in normal feeding without wings were: *Ubi* > + F (n=64), *Ubi* > + M (n=62), *Ubi* > *bmm-RNAi^V37877^* F (n=56), *Ubi* > *bmm-RNAi^V37877^* M (n=55), *Ubi > bmm-RNAi^V37880^* F (n=58) and *Ubi > bmm-RNAi^V37880^* M (n=61); “n” in starvation were: *Ubi* > + F (n=56), *Ubi* > + M (n=42), *Ubi* > *bmm-RNAi^V37877^* F (n=54), *Ubi* > *bmm-RNAi^V37877^* M (n=52), *Ubi > bmm-RNAi^V37880^* F (n=63) and *Ubi > bmm-RNAi^V37880^* M (n=63). Asterisks indicate significant differences at *p* < 0.05 compared with the appropriate female or male control, following Tukey’s correction for multiple tests. Error bars are SEM. ANOVA tests are reported in Table S2.

To verify that the effect on locomotor activity in *Ubi > bmm-RNAi^V37877^* males was not due to the spread-wings phenotype, we also recorded their activity after removing their wings. Although wing removal affected locomotor activity in all flies, the total 24h locomotor activity (Fig. 2d) and activity during the daytime (Fig. 2e) were still lowest in these males (see also Fig. 2f and Table S2). Total activity and daytime activity were also reduced in females of both *Ubi > bmm-RNAi* flies without wings compared to controls.

To assess the effect of starvation on locomotor activity when *bmm* expression is reduced, we recorded locomotor activity in DAM tubes containing 1% agar for 3 days. Flies that died during the assay were discarded from further analyses. All flies were more active under starvation conditions, but total 24h locomotor activity (Fig. 2g) and daytime activity (Fig. 2h) was still reduced in *Ubi > bmm-RNAi^V37877^* males and *Ubi > bmm-RNAi^V37880^* females (see also Fig. 2i and Table S2) compared with control flies. Interestingly, *Ubi > bmm-RNAi^V37880^* males were more active under food deprivation during the whole day and during the daytime (Fig. 2g and Fig. 2h).

### *bmm* downregulation results in altered sleep patterns

We next assessed whether impaired energy metabolism would affect sleep patterns in addition to locomotor activity (see ANOVA tests in Table S3). We evaluated sleep behavior of *Ubi* > *bmm-RNAi* flies and found that *Ubi > bmm-RNAi^V37880^* females had fewer sleep bouts than control flies (Fig. 3a). Conversely, males of both *bmm-RNAi* lines had more sleep bouts than control flies, but the bouts were shorter (Fig. 3b). *Ubi > bmm-RNAi^V37877^* males slept more during the daytime, but *Ubi > bmm-RNAi^V37880^* males slept less (Fig. 3c). In contrast, *Ubi > bmm-RNAi^V37877^* females had normal sleep bouts but rested more during the daytime. Sleep bouts were not affected in flies without wings (Fig. 3d), but *Ubi > bmm-RNAi^V37877^* males had longer sleep bouts (Fig. 3e) and slept more during the daytime and nighttime (Fig. 3f). Also, females with ubiquitous expression of both *bmm-RNAi* lines napped more during the daytime.

**Fig 3.**
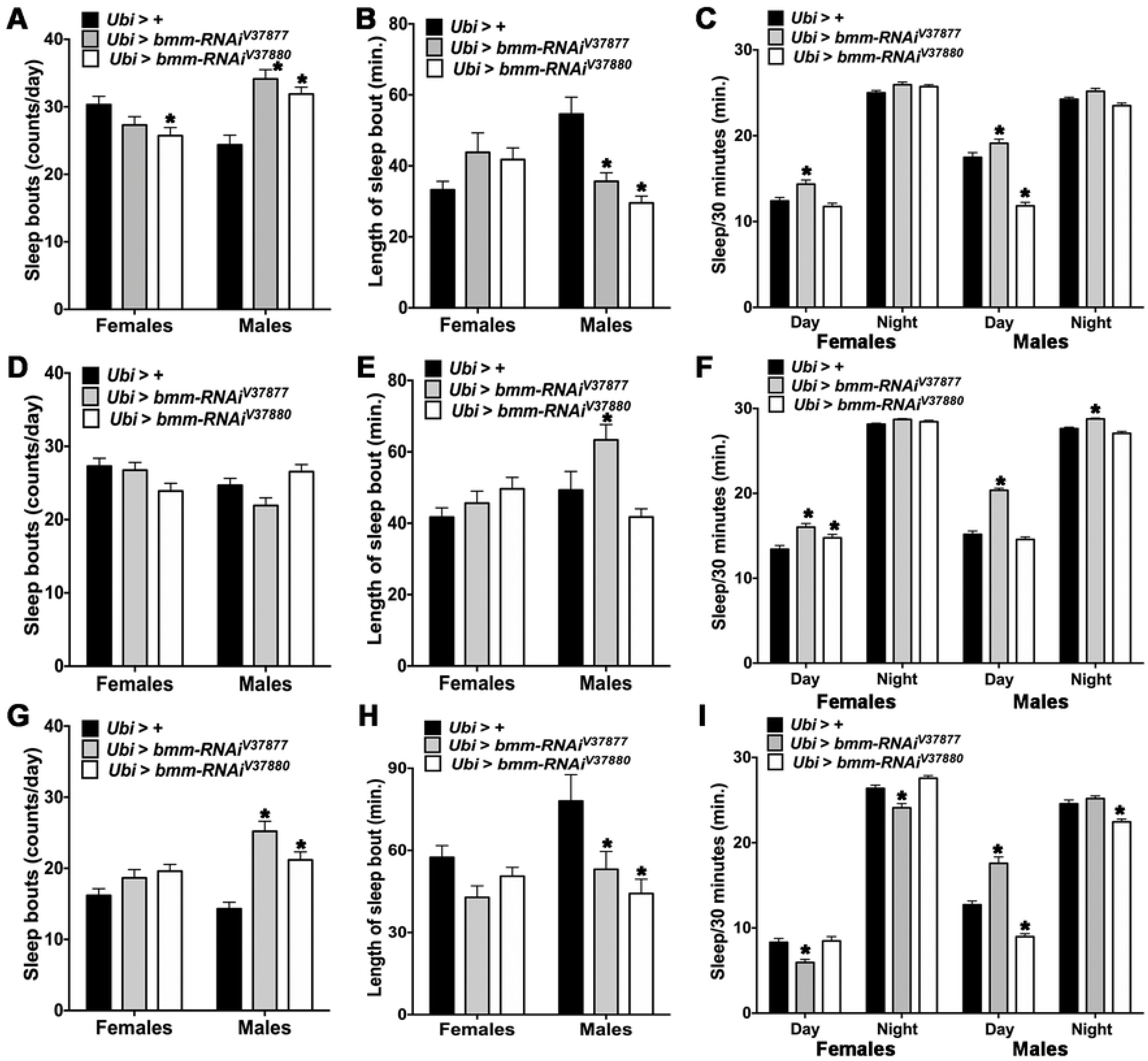
Sleep behavior in *Ubi* > *bmm-RNAi* flies. Offspring of *Ubi-Gal4* mated with GD control (*Ubi* > +) or *UAS-bmm-RNAi* flies (*Ubi* > *bmm-RNAi^V37877^* and *Ubi* > *bmm-RNAi^V37880^*) were used for the sleep assay in normal feeding (a-c), normal feeding without wings (d-f) and starvation conditions (g-i). (a, d and g) Average number of sleep bouts. (b, e and h) Average length of sleep bout. (c, f and i) Average sleep during the daytime and the nighttime. Sleep was calculated using the standard definition of a continuous period of inactivity lasting at least 5 minutes, and average was calculated from 7 days of behavior assay except for starvation, which was measured over 3 days, and recalculated to periods of 30 minutes of sleep. Asterisks indicate significant differences at *p* < 0.05 compared with the appropriate female or male control, following Tukey’s correction for multiple tests. Error bars are SEM. ANOVA tests are reported in Table S3.

Under starvation conditions all flies had less sleep bouts than under normal feeding (Fig. 3a and Fig. 3g). Starved *Ubi > bmm-RNAi* males of both lines had more sleep bouts (Fig. 3g) and the bouts were shorter (Fig. 3h) than starved control males. Interestingly, starved *Ubi > bmm-RNAi^V37877^* females and starved *Ubi > bmm-RNAi^V3880^* males slept less during the daytime and during the nighttime than the control (Fig. 3i). Starved *Ubi > bmm-RNAi^V37877^* males slept more during the daytime than the control (Fig. 3i). In contrast, starved *Ubi > bmm-RNAi^V37880^* males slept less during the daytime and nighttime than the corresponding control. Thus, effects of *bmm* expression on sleep patterns are condition-dependent and sex-dependent and vary with the extent of reduction in gene expression in the different RNAi lines.

### *bmm* downregulation in the fat body or oenocytes is sufficient to affect locomotor activity

Since the fat body and oenocytes are the two main tissues for triglyceride metabolism, we asked whether reduction of *bmm* expression specifically in these tissues could account for the observed effects on locomotor activity and sleep (Fig. S1 and S2, and Tables S4 and S5). We combined *UAS-bmm-RNAi* with *Lsp2-Gal4* to down-regulate *bmm* in the fat body, and with *Dsat1-Gal4* and *Ok72-Gal4* to down-regulate *bmm* in oenocytes. Females of both *Lsp2* > *bmm-RNAi* lines had higher total 24h locomotor activity than controls (Fig. S1a) and were more active during the daytime (Fig. S1b). In contrast, *Lsp2* > *bmm-RNAi^V37880^* males had lower 24h activity than controls (Fig. S1a) and were less active during the daytime (Figs. S1b-c). Interestingly, females of both *Dsat1* > *bmm-RNAi* lines showed less total 24h activity (Fig. S1d) and less daytime activity than control flies (Figs. S1e-f). This reduction in activity also occurred with *OK72* > *bmm-RNAi^V37880^* females (Figs. S1g-i). Furthermore, *Dsat1* > *bmm-RNAi^V37880^* males had lower 24h activity (Fig. S1d) and lower daytime activity than control flies (Fig. S1e). Similarly, *Dsat1* > *bmm-RNAi^V37877^* males exhibited reduced daytime activity (Fig. S1e).

### Tissue-specific reduction in *bmm* expression affects sleep behavior

*Lsp2 > bmm-RNAi* females of both strains had more sleep bouts (Fig. S2a) than control flies but their bouts were shorter (Fig. S2b) and these flies slept less during the daytime (Fig. S2c). Interestingly, females and males of both *bmm-RNAi* lines combined with the two oenocyte drivers were not affected in the number or length of sleep bouts (Figs. S2d-e and S2g-h); however, females slept more during the daytime (Figs. S2f and S2i). Moreover, *bmm-RNAi^V37880^* males combined with both oenocyte drivers slept more during the daytime similar to *Dsat1* > *bmm-RNAi^V37877^* males (Figs. S2f and S2i). Also, males of both *OK72* > *bmm-RNAi* flies rested less in the nighttime (Fig. S2i). Our results show that metabolic effects resulting from reduction of *bmm* expression in the fat body or oenocytes are sufficient to affect energy-dependent organismal phenotypes.

### *bmm* downregulation affects the metabolome in a sex-dependent manner

To gain insights in the mechanisms by which suppression of *bmm* expression results in physiological effects, we performed a comprehensive metabolomic analysis. The metabolome is a proximal link between gene expression and organismal phenotypes. To assess to what extent disruption of triglyceride metabolism through reduction of *bmm* expression affects the metabolome, we performed global metabolomics analysis on 6-day old *Ubi* > *bmm-RNAi* flies reared under standard culture conditions. We identified 767 metabolites, consisting of 723 known compounds and 44 compounds of unknown structural identity (Dataset S1). Principal components analysis (PCA) highlights sexual dimorphism of the metabolome, with males and females separated along Component 1 (31.1% of the variance between samples) and genotypes partially separated along Component 2 (12.4% of the variance) (Fig. 4a). Therefore, we performed all metabolomic analyses separately for females and males in single comparisons or comparing both *bmm-RNAi* lines with the corresponding control (Dataset S1). The metabolomes of *Ubi* > *bmm-RNAi^V37877^* females and males differ more from their controls (251 and 261 altered metabolite levels, respectively) than *Ubi* > *bmm-RNAi^V37880^* females and males (179 and 223 altered metabolite levels, respectively). However, there were quantitative changes in metabolites common to both genotypes: 124 in females and 118 in males (Fig. 4b and Fig 4d). The proportion of metabolites upregulated/downregulated was higher in *Ubi* > *bmm-RNAi^V37877^* (213↑/38↓ in females and 208↑/53↓ in males) than *Ubi* > *bmm-RNAi^V37880^* (101↑/78↓ in females and 94↑/129↓ in males) (Fig. 4c and Fig. 4e). These data reveal sex-dependent effects on the metabolome. The genotypes with higher effects on the metabolome correlate with lower levels of *bmm* transcripts (RT-qPCR), and greater effects on lifespan, locomotor activity and sleep parameters.

**Fig 4.**
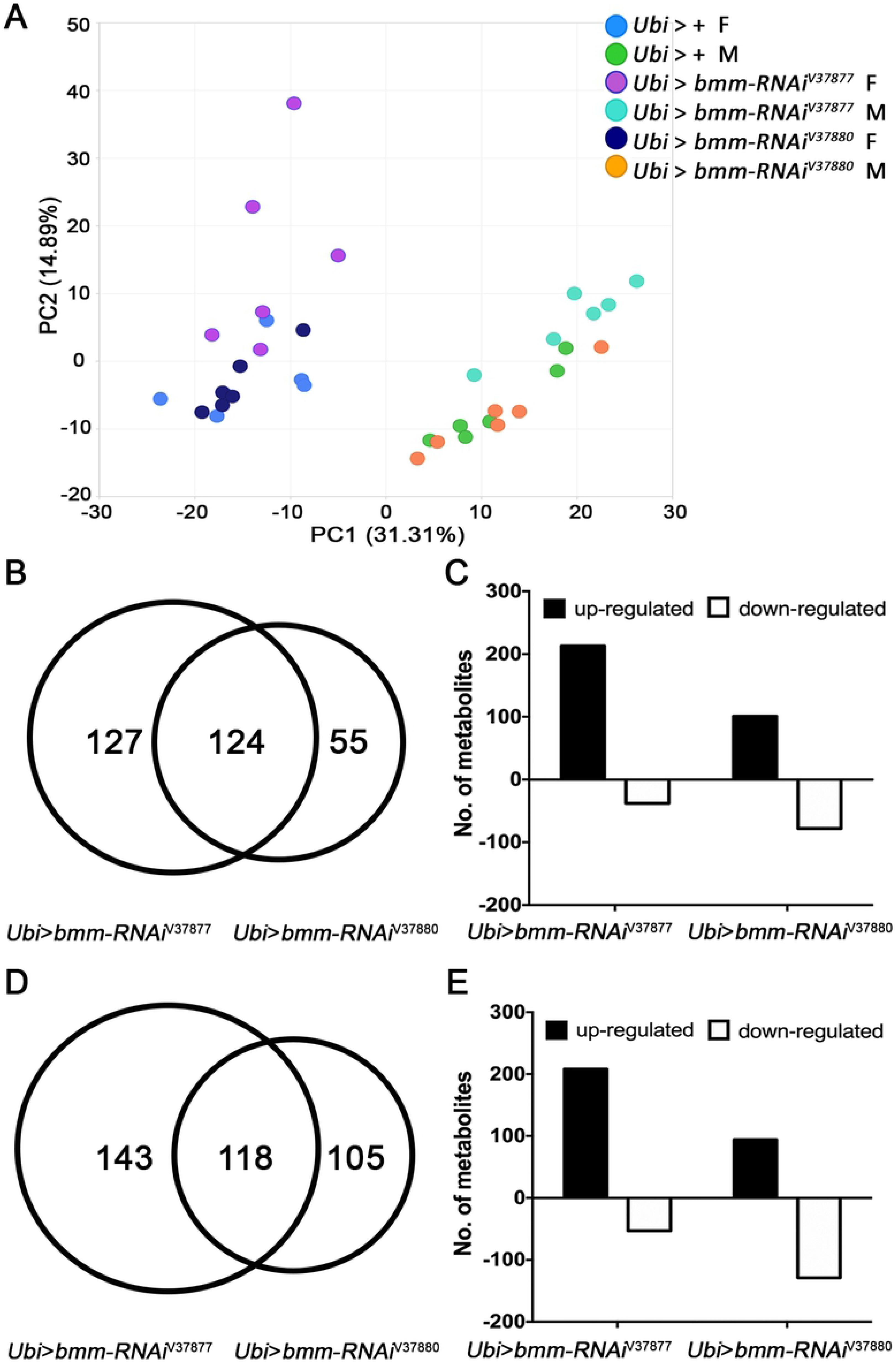
Metabolites with altered abundance levels in *Ubi* > *bmm-RNAi* flies. Offspring of *Ubi-Gal4* mated with GD control (*Ubi* > +) or *UAS-bmm-RNAi* flies (*Ubi* > *bmm-RNAi^V37877^* and *Ubi* > *bmm-RNAi^V37880^*) were used for metabolome analysis. F1 6-day-old adult females (F) and males (M) were used for the metabolomic profiles (n=6). ANOVA contrasts were used to identify metabolites that differed significantly between experimental groups and corresponding controls at *p* ≤ 0.05 (q-value is reported in Supplementary Dataset 1). (a) Principal component analysis of variation in the composition of the metabolome of flies in which *bmm* is downregulated and control flies. (b) Venn diagram of metabolites that significantly changed in *Ubi* > *bmm-RNAi* females compared with controls. (c) Number of metabolites that significantly increased (black bars) and decreased (white bars) in *Ubi* > *bmm-RNAi* females compared with controls. (d) Venn diagram of metabolites that significantly changed in *Ubi* > *bmm-RNAi* males compared with controls. (e) Number of metabolites that significantly increased (black bars) and decreased (white bars) in *Ubi* > *bmm-RNAi* males compared with controls.

We performed Metabolite Set Enrichment Analysis (MSEA) for differentially abundant metabolites of single comparisons (Dataset S2: a-b and d-e) or comparisons of both *bmm-RNAi* lines (Dataset S2: c and f) with corresponding controls. Overrepresentation analysis exhibited changes in metabolic pathways of lipids (e.g. “Fatty acid metabolism”), carbohydrates (e.g. “Glycogen metabolism” and “Glycolysis, gluconeogenesis and pyruvate metabolism”), amino acids (e.g. “Tryptophan metabolism” and “Glycine, serine and threonine metabolism”) and nucleotides (e.g. “Purine metabolism” and “Pyrimidine metabolism”) for all comparisons.

### *bmm* downregulation alters lipid metabolism

We analyzed 298 metabolites from 49 lipid metabolism sub-pathways in flies in which *bmm* expression was suppressed compared to controls (Fig. S3). *Ubi* > *bmm-RNAi^V37877^* females experienced more changes in metabolite levels (69, 57↑/12↓) than *Ubi* > *bmm-RNAi^V37880^* females (60, 25↑/35↓). Conversely, *Ubi* > *bmm-RNAi^V37877^* males had fewer changes in metabolite levels (76, 58↑/18↓) than *Ubi* > *bmm-RNAi^V37880^* males (125, 44↑/81↓).

Our metabolomic analysis revealed significant decreases in almost all free fatty acids and monoacylglycerols evaluated in *Ubi* > *bmm-RNAi^V37880^* males and in two lipid metabolites of *Ubi* > *bmm-RNAi^V37880^* females (Fig. 5). *Ubi* > *bmm-RNAi^V37877^* males also showed decreases in the same lipid species, but the relative decreases failed to meet the threshold of significance (Dataset S1). We also observed significant decreases in many acylcarnitine species in both sexes of *Ubi* > *bmm-RNAi^V37880^* (Fig. 6), indicating a deficiency in fatty acid transport into the mitochondria. These observations correlate with lipid enrichment analysis of differentially abundant metabolites which highlight “Fatty acyls” as the most prominent group (Dataset S3) in all comparisons versus control.

**Fig 5.**
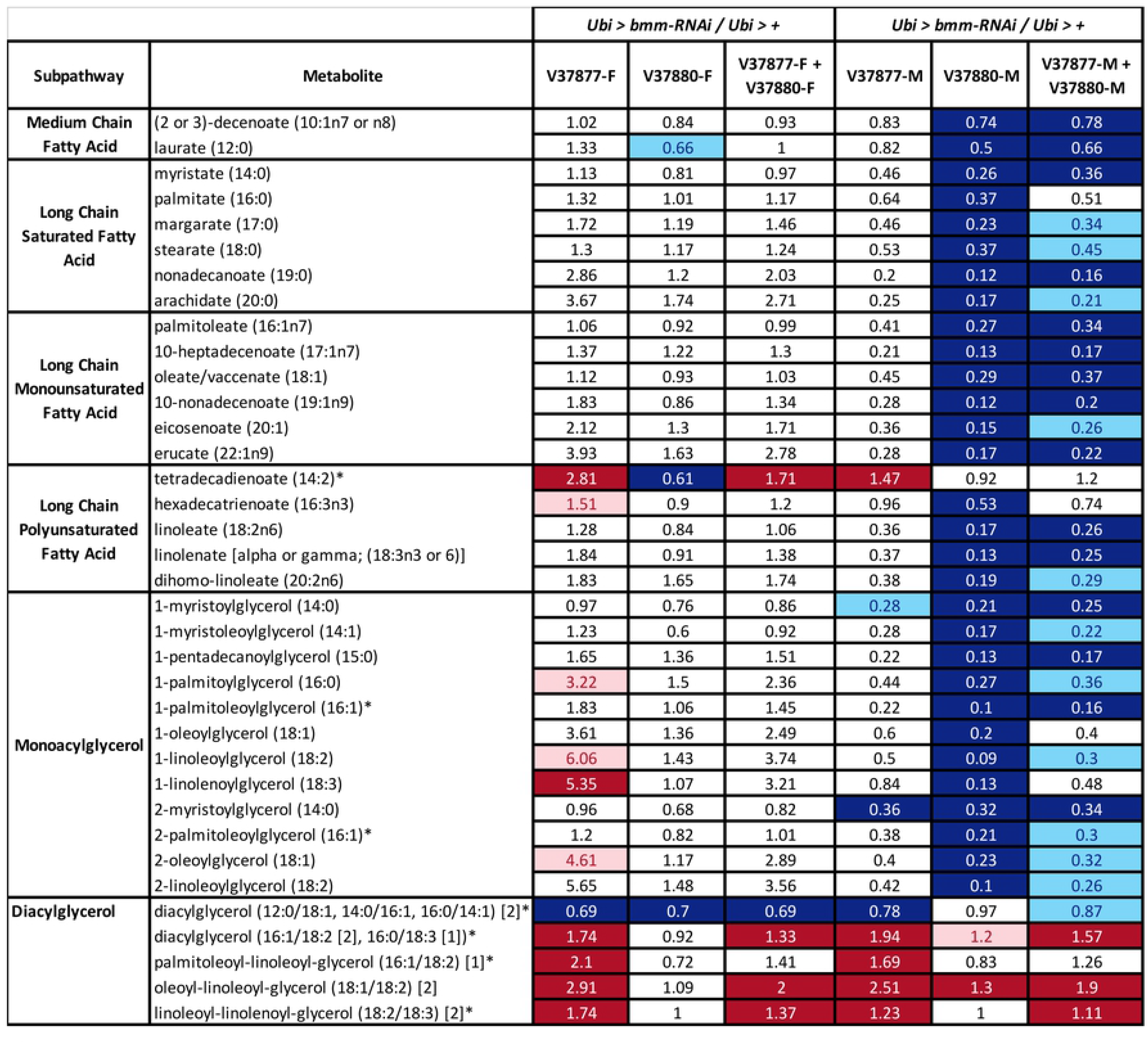
Compounds of lipid metabolism with altered levels of abundance in *Ubi* > *bmm-RNAi* flies. Heat map of changes in biochemicals of lipid metabolism in flies in which *bmm* is downregulated (*Ubi* > *bmm-RNAi*) compared with control flies (*Ubi* > +). For this graph and the graphs below, we performed a single comparison of each strain with its corresponding control for females (V37877-F and V37880-F) and males (V37877-M and V37880-M). Also, we performed a comparison of both strains with their corresponding controls for females (V37877-F + V37880-F) and males (V37877-M + V37880-M). Red and dark blue represent the metabolites that increased and decreased respectively at *p* ≤ 0.05, light red and light blue represent the metabolites that increased and decreased respectively at 0.05 ≤ *p* ≤ 0.1 (q-value is reported in Dataset S1). Asterisks indicate compounds that have not been confirmed based on a standard.

**Fig 6.**
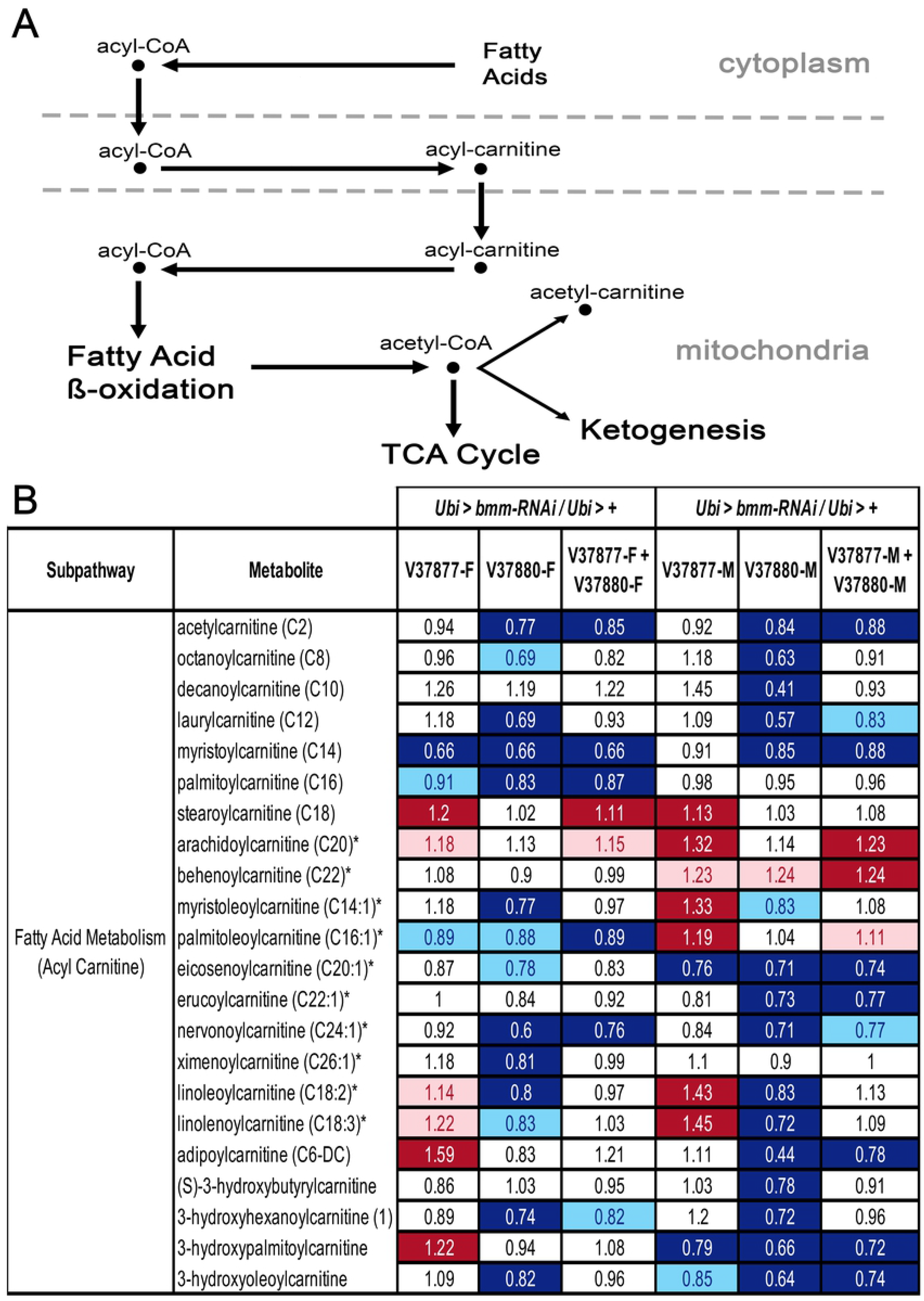
Compounds of acyl carnitine/fatty acid metabolism with altered abundance levels in *Ubi* > *bmm-RNAi* flies. (b) Diagram of the acyl carnitine/fatty acid metabolic pathway, highlighting the role of carnitine in facilitating transport of fatty acids into the mitochondria for fatty acid oxidation. (b) Heat map of changes in biochemicals of acyl carnitine/fatty acid metabolism in flies in which *bmm* is downregulated (*Ubi* > *bmm-RNAi*) compared with control flies (*Ubi* > +). Red and dark blue represent the metabolites that increased and decreased respectively at *p* ≤ 0.05, light red and light blue represent the metabolites that increased and decreased respectively at 0.05 ≤ *p* ≤ 0.1 (q-value is reported in Dataset S1). Asterisks indicate compounds that have not been confirmed based on a standard.

We observed significant increases in several unsaturated diacylglycerol species (Fig. 5), which suggests that conversion of triacylglycerol to diacylglycerol can occur in mutants with reduced expression of *bmm,* but that further breakdown of diacylglycerols is impaired. *Drosophila* utilizes a dual lipolytic strategy for fat mobilization, where Bmm seems to support the basal demands of lipolysis, while AkhR signaling, a cAMP-induced GPCR/PKA pathway, initiates the lipolytic response system that supports rapid fat mobilization [19, 27]. Indeed, we observed significantly higher levels of cAMP in *bmm* knock-down females (Dataset S1). Thus, the increase in long-chain diacylglycerols (DAGs), which are the most abundant DAGs in the fat body, brain and gut [28] in *Ubi* > *bmm-RNAi^V37877^* females and males could indicate that *bmm* knockdown causes a switch to a second lipolytic system to compensate for the energy demands when *bmm* is expressed at low levels during normal feeding. Whereas changes in diacylglycerol species are correlated for males and females, overall changes in abundances of other lipid metabolites are sexually dimorphic with females counteracting the reduction in *bmm* more efficiently (Fig. 5), as reported in other studies [25]. We also observed a significant increase in phosphatidylcholine (PC) and phosphatidylethanolamine (PE) levels (Dataset S1), suggesting increased activity of phospholipase C or increased *de novo* synthesis of PC and PE.

### Carbohydrate metabolism may compensate for lipid impairment in *bmm* knockdown flies

Impairment of lipid mobilization for energy production resulted in marked changes in the intermediates of the glycolysis pathway and the tricarboxylic acid (TCA) cycle in both *bmm* knockdown strains (Fig. 7 and Dataset S2). We evaluated 47 metabolites from 9 carbohydrate metabolism sub-pathways (Fig. S4). The same number of metabolites showed changes in abundance in *Ubi* > *bmm-RNAi^V37877^* females (15, 12↑/3↓) and in *Ubi* > *bmm-RNAi^V37880^* females (15, 10↑/5↓). However, more carbohydrate metabolites showed altered abundances in *Ubi* > *bmm-RNAi^V37877^* males (22, 19↑/3↓) than in *Ubi* > *bmm-RNAi^V37880^* males (11, 8↑/3↓). We observed more changes in metabolites associated with energy metabolism in *Ubi* > *bmm-RNAi^V37877^* females than any other genotypes (Fig. S5). Our data indicate that altered carbohydrate metabolism in the *bmm* knockdown strains could compensate for impaired lipid mobilization.

**Fig 7.**
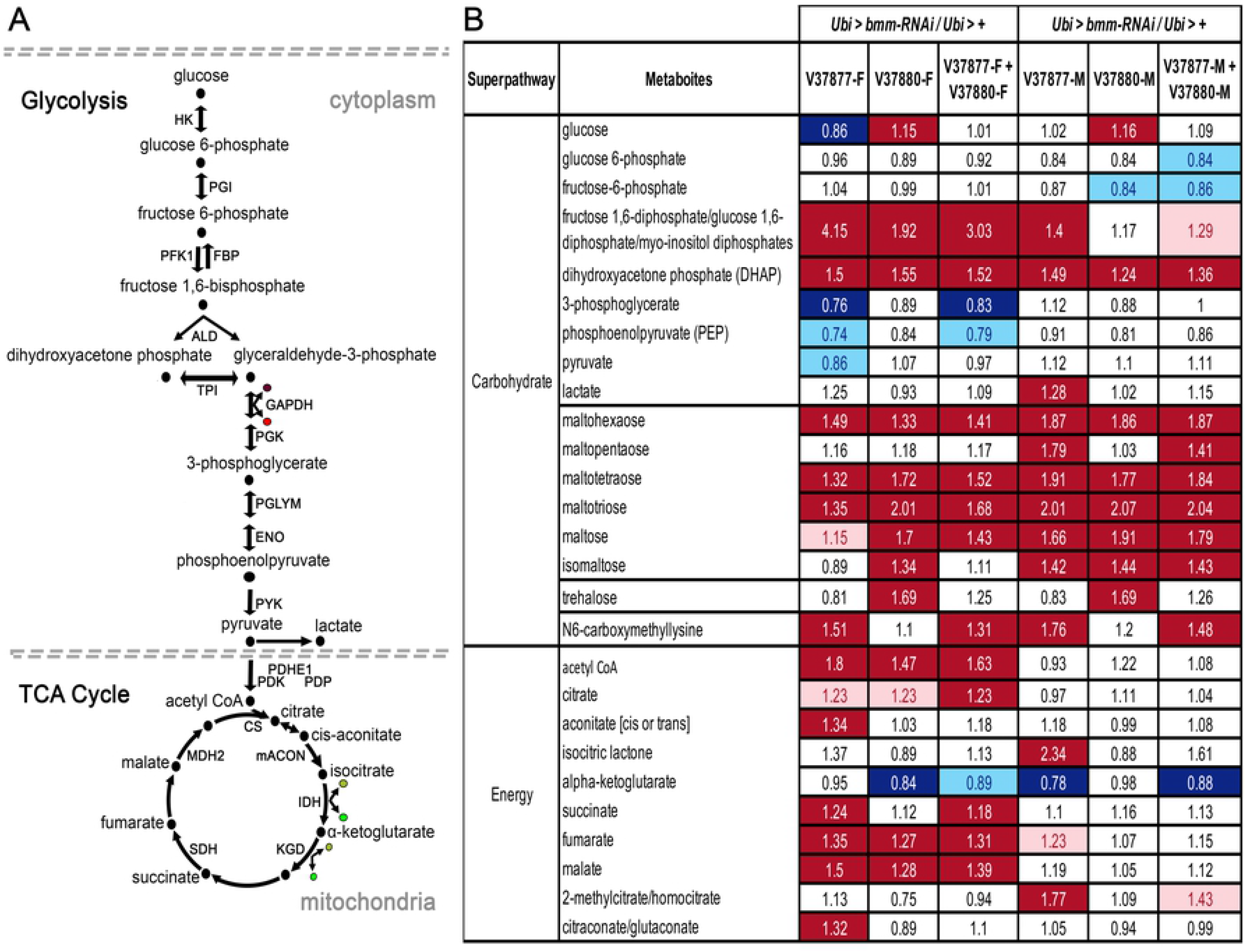
Metabolites of glycolysis and the TCA cycle with altered abundance levels in *Ubi* > *bmm-RNAi* flies. (a) Diagram of the glycolysis pathway and the TCA cycle. Black dots represent intermediates in the pathways and the enzymes in the reactions are marked with arrows and their initials: HK=Hexokinase, PGI=Phosphoglucose isomerase, FBP=Fructose 1,6-biphosphatase, PFK=Phosphofructokinase, ALD=Aldolase, TPI=Triosephosphate isomerase, GAPDH=Glyceraldehyde-3-phosphate dehydrogenase, PGK=Phosphoglycerate kinase, PGLYM=Phosphoglyceromutase, ENO=Enolase, PYK=Pyruvate kinase, PDHE1=Pyruvate dehydrogenase E1, PDK=Pyruvate dehydrogenase kinase, PDP=Pyruvate dehydrogenase phosphatase, CS=Citrate synthase, mACON=Mitochondrial aconitase, IDH=Isocitrate dehydrogenase, KGD=alpha-ketoglutarate dehydrogenase, SDH=Succinate dehydrogenase, MDH2=Malate dehydrogenase. Brown dots designate ADP, bright-red dots, ATP, opaque-green dots, NAD+ and bright-green dots, NADH. (b) Heat map of variation in intermediates of carbohydrate metabolism and TCA cycle with altered abundance levels in *bmm* down-regulated (*Ubi* > *bmm-RNAi*) and control flies (*Ubi* > +). Red and dark blue represent the metabolites that increased and decreased respectively at *p* ≤ 0.05, light red and light blue represent the metabolites that increased and decreased respectively at 0.05 ≤ *p* ≤ 0.1 (q-value is reported in Dataset S1). Asterisks indicate compounds that have not been confirmed based on a standard.

### *bmm* downregulation results in changes in amino acid and nucleotide metabolic pathways

The shifts in energy metabolism described above are accompanied by changes in intermediaries of amino acid and peptide metabolism pathways (Fig. S6 and S7, and Dataset S2). We evaluated 175 compounds in 14 amino acid metabolic pathways. *Ubi* > *bmm-RNAi^V37877^* females had altered quantities of 65 metabolites (58↑/7↓), while only 39 metabolites changed (27↑/12↓) in *Ubi* > *bmm-RNAi^V37880^* females. *Ubi* > *bmm-RNAi^V37877^* males showed changes in 64 metabolites (54↑/10↓), while *Ubi* > *bmm-RNAiV^37880^* males only showed alterations in 32 compounds (18↑/14↓).

We also identified 76 metabolites from 9 nucleotide metabolism sub-pathways (Fig. S8). *Ubi* > *bmm-RNAi^V37877^* flies showed more changes in levels of nucleotide intermediaries than *Ubi* > *bmm-RNAi^V37880^* flies. In *Ubi* > *bmm-RNAi^V37877^* females 65 metabolites underwent changes in abundance (58↑/7↓), whereas only 39 metabolites changed (27↑/12↓) in *Ubi* > *bmm-RNAi^V37880^* females. In *Ubi* > *bmm-RNAi^V37877^* males 64 metabolites showed altered levels of abundance (54↑/10↓) and levels of 32 metabolites changed in *Ubi* > *bmm-RNAi^V37880^* males (18↑/14↓). Thus, dysregulation of lipid metabolism has widespread consequences in the metabolome.

### *bmm* knockdown alters neurotransmitter levels

Motivated by the effects of *bmm* suppression on locomotion, sleep and lifespan, we explored changes in neurotransmitter levels resulting from *bmm* knockdown. We observed significant alterations, dependent on RNAi genotype and sex, of different neurotransmitters in *bmm* knockdown flies (Dataset S1). Inhibition of the tryptophan-kynurenine (TRY-KYN) pathway increases N-acetylserotonin (NAS) levels, which has been associated with extended lifespan in *Drosophila* [29, 30]. Indeed, we found reduced levels of NAS in both sexes of *bmm* knockdown flies and elevated TRY-KYN pathway metabolites in line with their reduced lifespan (Fig. 1C). In addition, levels of serotonin were increased in *bmm* knockdown males. Furthermore, dopamine and its precursor l-DOPA are elevated in *bmm* knockdown flies. We also found altered levels of tyramine in *bmm* knockdown females and of acetylcholine in both sexes.

## Discussion

We used RNAi-mediated suppression of *bmm*, which encodes triacylglyceride lipase in *Drosophila*, to explore the behavioral, physiological and metabolic consequences of disruption of lipid mobilization. We mitigated potential off-target effects using two different RNAi lines targeting *bmm* with no predicted off-target effects. Both RNAi constructs affected the same phenotypes, but with quantitative differences that largely correlated with their different efficacies in *bmm* suppression. We observed pervasive sex differences in all organismal level phenotypes and in the metabolome, which is not surprising as sexual dimorphism has been reported for virtually all complex traits examined in *Drosophila* [15,31–33]. Reduction in expression of *bmm* affected locomotor activity and sleep patterns and resulted in widespread shifts in abundances of metabolites in a range of metabolic pathways. We note, however, that the metabolomic data were obtained at a single time point during the circadian cycle and at a single age. Sampling at different times during the day and night and at different ages may provide more comprehensive information on the circadian dynamics of the effects of *bmm* suppression on the metabolome.

A previous study showed that a restricted larval diet results in downregulation of *bmm* [34]. Adults that emerged from these larvae had increased levels of triglycerides and were more resistant to starvation stress. This effect could be mimicked by targeted reduction of *bmm* levels in the larval fat body [34]. Reduction of *bmm* expression in the fat body in adult flies also protects against cardiac dysfunction during starvation and flies with reduced *bmm* expression during starvation have higher levels of metabolites involved in energy metabolism, and high levels of diacylglycerol [35], which is also upregulated under our experimental conditions (Fig 5).

### Tissue-specific inhibition of *bmm* in the fat body and oenocytes is sufficient to affect locomotor activity and sleep patterns

Our locomotor activity data show the strongest effect on locomotor activity both under normal feeding conditions and during food deprivation in *Ubi > bmm-RNAi^V37877^* males, consistent with the lowest level of *bmm* transcripts in these flies (Fig. 1 and 2). Furthermore, *bmm* downregulation in fat body or oenocytes was sufficient to affect locomotor activity and sleep (Fig. S1 and S2). In females, however, effects on locomotor activity are in opposite directions when *bmm* is downregulated in the fat body and oenocytes. *bmm* function in the fat body is necessary for lipid utilization by oenocytes, and oenocytes are essential for lipid regulation, but the midgut can also provide TAGs to oenocytes [18]; thus, females might use another source of lipids when *bmm* is downregulated in fat body to compensate. These data suggest that *bmm* function in oenocytes is a limiting factor to obtain energy for locomotor activity in both sexes, but *bmm* function in the fat body is more important in males, such as it is in the somatic cells of the gonads [25].

The effect of *bmm* suppression in the fat body and oenocytes on locomotion and sleep was weaker than the effect observed with a ubiquitous *Gal4* driver. This could be due to less effective suppression of *bmm* with these tissue-specific drivers or to intact lipolysis in other tissues that may contribute energy towards locomotion and normal sleep patterns. *bmm* is expressed in different tissues [23]; indeed, neuronal function of *bmm* is necessary to regulate sex-dependent triglyceride breakdown [25]. Sleep disruption is associated with increased weight and lipids in humans [36, 37] but it is not clear if dysregulation of lipids affects sleep [38].

### Carbohydrate metabolism may compensate for lipid impairment in *bmm* knockdown flies

We observed significant increases in carbohydrates, such as maltohexaose and maltopentaose, as well as glucose and trehalose levels in *Ubi* > *bmm-RNAi^V37880^* flies. Trehalose is a disaccharide consisting of two glucose molecules, which serves as a source for glucose in the hemolymph [39]. Since *Ubi* > *bmm-RNAi^V37880^* flies exhibited a severely impaired lipid mobilization phenotype, the fat body in these flies may generate trehalose to support energy demands and buffer the levels of glucose. Alternatively, decreased uptake and utilization of trehalose from the circulation could account for its accumulation in flies in which *bmm* expression is suppressed.

*Ubi* > *bmm-RNAi^V37877^* flies showed a significant increase in the advanced-glycation end product (AGE) N6-carboxymethyllysine, which serves as a biomarker of high blood glucose levels in humans; this might suggest a diabetes-like phenotype. High levels of AGEs together with high-fat diet produces liver dysfunction in mice [40], and liver dysfunction produces AGEs accumulation in humans [41]. Whether AGEs represent dysfunction of the fat body in *Ubi* > *bmm-RNAi^V37877^* flies is not clear, since glucose did not increase significantly in these flies, nor did they exhibit an increase in trehalose production by the fat body. These results provide a framework for future studies on compensatory metabolic mechanisms through manipulation of carbohydrate metabolism in the *bmm* mutant.

### *bmm* knockdown alters neurotransmitter levels

Inhibition of *bmm* expression results in elevated levels of the neurotransmitters serotonin and dopamine, which may account for impairment of locomotor activity and altered sleep patterns [42–44]. Serotonin 5-HT_1A_ receptors in *Drosophila* are required for insulin-producing cells to regulate lipid content [45]. Interestingly, the quantity of serotonin transporters (SERT) in the midbrain correlates with body-mass index in humans, with a negative correlation in people with obesity and a positive correlation in non-obesity individuals. This correlation supports the idea that serotonin may play a role in the reward system of food intake [46]. Also, increased levels of dopamine can lead to a reduction in striatal D2-dopamine receptors in humans with obesity through habituation [47]. Causal relationships between neurotransmitter levels and *bmm*-sensitive phenotypes in flies requires assessment in future experiments.

## Conclusions

Impairment of fatty acid mobilization results in a shift in intermediary metabolism toward utilization of carbohydrates as an energy source. In addition, amino acid, nucleotide, and neurotransmitter metabolic pathways are affected by disruption of lipid catabolism. This altered metabolic state gives rise to changes in morphological and physiological organismal phenotypes. Because pathways of intermediary metabolism are conserved across phyla, studies on the *Drosophila* model are relevant to advancing our understanding of human metabolic disorders.

## Materials and methods

### Fly stocks

We obtained two *w^1118^; UAS-bmm-RNAi* lines (#37877 and #37880) and their GD control from the Vienna *Drosophila* Resource Center [48]. All *Gal4* driver lines were obtained from the Bloomington *Drosophila* Stock Center: *w\**; *Ubi-Gal4*/*CyO* (#32551) for ubiquitous expression, *w^1118^; Lsp2-Gal4* (#6357) for specific expression in fat body, and *w*; Dsat1-Gal4* (#65405) and *w\**; *OK72-Gal4* (#6486) for expression in oenocytes. Flies were reared on molasses-cornmeal medium (Nutri-Fly®) with propionic acid and Tegosept (Genesee Scientific, Inc.) added as fungicides. They were maintained at 25°C and a 12h:12h light-dark schedule (lights on at 6:00 am). Matings were performed with 6 pairs per vial and flies were maintained at a controlled population density. *UAS-bmm-RNAi* flies and control GD flies were crossed with *Ubi-Gal4*/*CyO*. The progeny with one copy of *Ubi-Gal4* and *UAS-bmm-RNAi* (*Ubi > bmm-RNAi^V37877^* and *Ubi > bmm-RNAi^V37880^*) or *Ubi-Gal4* and GD control background (*Ubi > +*), were subjected to RT-qPCR, lifespan assay, locomotor activity/sleep assay and metabolomic analysis.

### RT-qPCR

Three biological replicates of 3 to 5-day old virgin females and males, with one copy of *Ubi-Gal4* and *UAS-bmm-RNAi* (*Ubi* > *bmm-RNAi^V37877^* and *Ubi > bmm-RNAi^V37880^*) or the GD control (*Ubi* > +), were anesthetized with CO_2_ and frozen on dry ice. All flies were collected at the same time of day to avoid effects of circadian rhythms and stored at -80°C. Total RNA was extracted using the RNeasy Plus Mini Kit (Qiagen, Inc.) and single strand cDNA was generated with High-Capacity cDNA Reverse Transcription (Thermo Fisher, AB). We designed primers to generate a PCR fragment of 116 bp for *bmm* and 102 bp for *Gapdh1* (Dataset S4). Relative quantification (RQ) was done using the 2^-ΔΔCt^ method with *Gapdh1* as internal control because it was stable across genotypes when comparing raw cycle thresholds (Tukey’s test p < 0.05, n=9). Three technical replicates were used for each sample, and qPCR was performed in a QuantStudio 3 instrument (Applied Biosystems).

### Lifespan assay

We used mated flies with one copy of *Ubi-Gal4* and *UAS-bmm-RNAi* (*Ubi* > *bmm-RNAi^V37877^* and *Ubi > bmm-RNAi^V37880^*) or the GD control (*Ubi* > +) for lifespan assays. We set up 50 vials of molasses-cornmeal media with 3 males or 3 females for each genotype, transferred them to fresh food every 2-3 days, and scored for survival daily. Flies that escaped during transfers or were stuck in the food were discarded from the analysis (Dataset S5). Average lifespan for each group was calculated and data are presented as mean ± standard error.

### Locomotor activity/sleep assay

*w^1118^; UAS-bmm-RNAi* flies and control GD flies were crossed with the ubiquitous driver (*w*; Ubi-Gal4*/*CyO*), the fat body driver (*w^1118^; Lsp2-Gal4*) or oenocyte drivers (*w*; OK72-Gal4* and *w*; Dsat1-Gal4*). Offspring with one copy of driver-*Gal4* and *UAS-bmm-RNAi* or the GD control were subjected to activity monitoring using *Drosophila* Activity Monitors (DAMSystem3, TriKinetics Inc.), which record movement by counting interruptions of an infrared beam. Tubes were plugged with 2% agar and 5% sucrose for normal feeding assays and 1% agar for starvation assays [31].

Two-day old males and females of the three genotypes were collected simultaneously and unmated flies were used for the assay. Two full 32-tube replicates were analyzed for activity behavior for each sex and genotype (Datasets S6-S10). The incubator followed a 12h day-12h night cycle and the activity behavior was recorded for 7 days in one-minute bins. Whole day activity, daytime/nighttime activity, average activity profiles, sleep bouts, sleep bout length and actograms were calculated using ShinyR-DAM^49^ and data are presented as mean ± standard error. Means were calculated from 7 days of recording except for starvation where only 3 days were recorded. Sleep was calculated using the standard definition of a continuous period of inactivity lasting at least 5 min [50]. Sleep events were identified using a sliding window algorithm of 5 min width and 1 min sliding interval, then averaged over all individual flies per condition.

### Metabolomic analysis

Six replicates of ∼150 6-day old females and males for each genotype (36 samples) were collected at the same time of day on dry ice and stored at -80°C. Unmated flies were used for metabolomic profiling by Metabolon, Inc., NC. Samples were prepared using the automated MicroLab STAR system from Hamilton Company, adding several recovery standards for quality control prior to the extraction process. Proteins were precipitated with methanol under vigorous shaking (Glen Mills GenoGrinder 2000) for 2 min., followed by centrifugation (to remove protein, dissociate small molecules bound to protein or trapped in the precipitated protein matrix, and to recover chemically diverse metabolites). The extract was divided into five fractions: two for analysis by two separate RP/UPLC-MS/MS methods with positive ion mode ESI, one for analysis by RP/UPLC-MS/MS with negative ion mode ESI, one for analysis by HILIC/UPLC-MS/MS with negative ion mode ESI, and one sample was reserved for backup. Organic solvent was removed placing samples on a TurboVap (Zymark) and the sample extracts were stored overnight under nitrogen before preparation for analysis. Metabolites were extracted from the flies and loaded in an equivalent manner across the analytical platforms and the values for each metabolite were normalized based on Bradford protein concentrations.

Metabolon’s hardware and software were used for raw data extraction, peak-identification, and quality control processing. Compounds were identified by comparison to library entries of purified standards (or recorded as unknown entities) and peaks were quantified using area under the curve. Data normalization was performed to correct variation from instrument inter-day tuning differences. Each compound was corrected in run day blocks by registering the medians to equal one (1.00) and normalizing each data point proportionately. The detailed procedure for metabolomic profiling from Metabolon Inc. has been described previously [15].

### Metabolic networks and Metabolite Set Enrichment Analysis

Metabolic networks are rendered using a private custom software application made by Metabolon that is based on the Cytoscape application. This software uses the Cytoscape JS toolkit and the D3 JavaScript library [51, 52] and is available in MyMetabolon portal. Metabolite set enrichment analysis of global metabolic pathways (Supplementary Dataset 2) was performed with Fisher’s exact tests, mapping the list of differentially abundant metabolites to the Metabolon library, and the FDR method was used for correction of multiple comparisons. Enrichment analysis for lipid chemical structure (Dataset S3) was done using the PubChem-IDs lists as input in the “Enrichment Analysis” web tool of MetaboAnalyst 5.0 (https://www.metaboanalyst.ca). Mapped metabolites in the Human Metabolome Database (HMDB) were used for over-representation analysis of lipid structure comparing to the super-class dataset.

### Statistical analyses

Two-way ANOVA tests were performed for RT-qPCR and locomotor activity/sleep, followed by Tukey’s multiple testing correction (p<0.05) to assess statistically significant results (Tables S1-S5). For lifespan, we performed a two-way mixed model ANOVA with genotype and sex as fixed effects and vial as random effect, followed by Dunnett’s correction for multiple tests (p<0.05) (Supplementary Table S1). Metabolites profiled by mass spectroscopy in *Ubi > bmm-RNAi* females and males were analyzed separately with one-way ANOVA. ANOVA contrasts between experimental groups and controls were performed using two approaches: single comparisons of each strain with the corresponding control (V37877-F, V37880-F, V37877-M and V37880-M) or comparisons of both strains with the proper control (V37877-F + V37880-F and V37877-M + V37880-M). Mean, p-values and q-values, and the magnitude of changes are in Dataset S1. Statistics and graphs were built on Prism 6 (GraphPad Software), R (R 3.6.1) and JMP (SAS Institute Inc.).

### Data Availability

*UAS-bmm-RNAi* lines and their GD control are available at the Vienna *Drosophila* Resource Center: https://stockcenter.vdrc.at/control/library_rnai and all *Gal4* driver lines are available from the Bloomington *Drosophila* Stock Center: https://bdsc.indiana.edu. The datasets supporting the conclusions of this article are included within the article and its supporting information. Metabolites profiled by mass spectroscopy in *Ubi > bmm-RNAi* flies, and statistics of metabolites that differed significantly between experimental groups and controls are in Dataset S1. MSEA data for global metabolic pathways and lipid structure of differentially abundant metabolites are in Dataset S2 and S3. The detailed procedure for metabolomic profiling from Metabolon Inc. has been described previously by Zhou *et al.* 2020 [15] at https://genome.cshlp.org/content/30/3/392/suppl/DC1. Our data of RT-qPCR: raw cycle thresholds and relative quantification (RQ) of *Gapdh* and *bmm* are in Dataset S4. Two factor ANOVA tests for RT-qPCR, lifespan and locomotor activity/sleep are in Tables S1-S5. All raw data and means are provided in Datasets 5-9.

## Acknowledgments

We would like to thank Dr. Therese A. Markow for comments on the manuscript

## Supporting information

**Fig S1. Locomotor activity in *bmm-RNAi* lines expressed in fat-body and oenocytes.** GD control and *UAS-bmm-RNAi* flies were mated with *Lsp2-Gal4* (a-c), *Dsat1-Gal4* (d-f) and *OK72-Gal4* (g-i), and F1-flies were used for the assay in normal feeding. (a, d and g) Average of whole-day locomotor activity. (b, e and h) Average of locomotor activity during the daytime and nighttime. (c, f and i) Average activity profiles in Zeitgeber time. Averages were calculated from 7 days of behavior assay. For this and Supplementary Fig. S2, “n” for females (F) and males (M) were: *Lsp2* > + F (n=60), *Lsp2* > + M (n=63), *Lsp2* > *bmm-RNAi^V37877^* F (n=55), *Lsp2* > *bmm-RNAi^V37877^* M (n=57), *Lsp2 > bmm-RNAi^V37880^* F (n=57), *Lsp2 > bmm-RNAi^V37880^* M (n=61), *Dsat1* > + F (n=62), *Dsat1* > + M (n=63), *Dsat1* > *bmm-RNAi^V37877^* F (n=63), *Dsat1* > *bmm-RNAi^V37877^* M (n=63), *Dsat1 > bmm-RNAi^V37880^* F (n=63), *Dsat1 > bmm-RNAi^V37880^* M (n=63), *OK72* > + F (n=58), *OK72* > + M (n=54), *OK72* > *bmm-RNAi^V37877^* F (n=61), *OK72* > *bmm-RNAi^V37877^* M (n=60), *OK72 > bmm-RNAi^V37880^* F (n=63) and *OK72 > bmm-RNAi^V37880^* M (n=61). Asterisks indicate significant differences at *p* < 0.05 compared with the appropriate female or male control, following Tukey’s correction for multiple tests. Error bars are SEM. ANOVA tests are reported in Table S4.

**Fig S2. Sleep behavior in *bmm-RNAi* lines expressed in fat-body and oenocytes.** GD control and *UAS-bmm-RNAi* flies were mated with *Lsp2-Gal4* (a-c), *Dsat1-Gal4* (d-f) and *OK72-Gal4* (g-i), and F1-flies were used for the assay in normal feeding. (a, d and g) Average number of sleep bouts. (b, e and h) Average length of sleep bout. (c, f and i) Average sleep during the daytime and the nighttime. Sleep was calculated using the standard definition of continuous period of inactivity lasting at least 5 minutes. Averages were calculated from 7 days of behavior assay and recalculated to periods of 30 minutes of sleep. Asterisks indicate significant differences at *p* < 0.05 compared with the appropriate female or male control, following Tukey’s correction for multiple tests. Error bars are SEM. ANOVA tests are reported in Table S5.

**Fig S3. Lipid metabolism network with biochemicals with significantly altered abundances in *Ubi* > *bmm-RNAi* flies.** (a) Metabolites that significantly changed in *Ubi* > *bmm-RNAi^V37877^* females compared with control females. (b) Metabolites that significantly changed in *Ubi* > *bmm-RNAi^V37880^* females compared with control females. (c) Metabolites that significantly changed in *Ubi* > *bmm-RNAi^V37877^* males compared with control males. (d) Metabolites that significantly changed in *Ubi* > *bmm-RNAi^V37880^* males compared with control males. Yellow nodes represent the sub-pathways analyzed (see Supplementary Dataset 1). Red and dark blue represent the metabolites that increased and decreased respectively at *p* ≤ 0.05, light red and light blue represent the metabolites that increased and decreased respectively at 0.05 ≤ *p* ≤ 0.1, and size of circles represent the magnitude of change. ANOVA contrasts were used to identify metabolites that differed significantly between experimental groups and proper controls, and q-value is reported in Dataset S1.

**Fig S4. Carbohydrate metabolism network with biochemicals with significantly altered abundances in *Ubi* > *bmm-RNAi* flies.** (a) Metabolites that significantly changed in *Ubi* > *bmm-RNAi^V37877^* females compared with control females. (b) Metabolites that significantly changed in *Ubi* > *bmm-RNAi^V37880^* females compared with control females. (c) Metabolites that significantly changed in *Ubi* > *bmm-RNAi^V37877^* males compared with control males. (d) Metabolites that significantly changed in *Ubi* > *bmm-RNAi^V37880^* males compared with control males. Yellow nodes represent the sub-pathways analyzed (see Supplementary Dataset 1). Red and dark blue represent the metabolites that increased and decreased respectively at *p* ≤ 0.05, light red and light blue represent the metabolites that increased and decreased respectively at 0.05 ≤ *p* ≤ 0.1, and size of circles represent the magnitude of change. ANOVA contrasts were used to identify metabolites that differed significantly between experimental groups and proper controls, and q-value is reported in Dataset S1.

**Fig S5. Network of energy metabolites with significantly altered abundances in *Ubi* > *bmm-RNAi* flies.** (a) Metabolites that significantly changed in *Ubi* > *bmm-RNAi^V37877^* females compared with control females. (b) Metabolites that significantly changed in *Ubi* > *bmm-RNAi^V37880^* females compared with control females. (c) Metabolites that significantly changed in *Ubi* > *bmm-RNAi^V37877^* males compared with control males. (d) Metabolites that significantly changed in *Ubi* > *bmm-RNAi^V37880^* males compared with control males. Yellow nodes represent the sub-pathways analyzed (see Supplementary Dataset 1). Red and dark blue represent the metabolites that increased and decreased respectively at *p* ≤ 0.05, light red and light blue represent the metabolites that increased and decreased respectively at 0.05 ≤ *p* ≤ 0.1, and size of circles represent the magnitude of change. ANOVA contrasts were used to identify metabolites that differed significantly between experimental groups and proper controls, and q-value is reported in Dataset S1.

**Fig S6. Amino acid metabolism network with intermediates with significantly altered abundances in *Ubi* > *bmm-RNAi* flies.** (a) Metabolites that significantly changed in *Ubi* > *bmm-RNAi^V37877^* females compared with control females. (b) Metabolites that significantly changed in *Ubi* > *bmm-RNAi^V37880^* females compared with control females. (c) Metabolites that significantly changed in *Ubi* > *bmm-RNAi^V37877^* males compared with control males. (d) Metabolites that significantly changed in *Ubi* > *bmm-RNAi^V37880^* males compared with control males. Yellow nodes represent the sub-pathways analyzed (see Supplementary Dataset 1). Red and dark blue represent the metabolites that increased and decreased respectively at *p* ≤ 0.05, light red and light blue represent the metabolites that increased and decreased respectively at 0.05 ≤ *p* ≤ 0.1, and size of circles represent the magnitude of change. ANOVA contrasts were used to identify metabolites that differed significantly between experimental groups and proper controls, and q-value is reported in Dataset S1.

**Fig S7. Peptide metabolism network intermediates with significantly altered abundances in *Ubi* > *bmm-RNAi* flies.** (a) Metabolites that significantly changed in *Ubi* > *bmm-RNAi^V37877^* females compared with control females. (b) Metabolites that significantly changed in *Ubi* > *bmm-RNAi^V37880^* females compared with control females. (c) Metabolites that significantly changed in *Ubi* > *bmm-RNAi^V37877^* males compared with control males. (d) Metabolites that significantly changed in *Ubi* > *bmm-RNAi^V37880^* males compared with control males. Yellow nodes represent the sub-pathways analyzed (see Dataset S1). Red and dark blue represent the metabolites that increased and decreased respectively at *p* ≤ 0.05, light red and light blue represent the metabolites that increased and decreased respectively at 0.05 ≤ *p* ≤ 0.1, and size of circles represent the magnitude of change. ANOVA contrasts were used to identify metabolites that differed significantly between experimental groups and proper controls, and q-value is reported in Dataset S1.

**Fig S8. Nucleotide metabolism network intermediates with significantly altered abundances in *Ubi* > *bmm-RNAi* flies.** (a) Metabolites that significantly changed in *Ubi* > *bmm-RNAi^V37877^* females compared with control females. (b) Metabolites that significantly changed in *Ubi* > *bmm-RNAi^V37880^* females compared with control females. (c) Metabolites that significantly changed in *Ubi* > *bmm-RNAi^V37877^* males compared with control males. (d) Metabolites that significantly changed in *Ubi* > *bmm-RNAi^V37880^* males compared with control males. Yellow nodes represent the sub-pathways analyzed (see Dataset S1). Red and dark blue represent the metabolites that increased and decreased respectively at *p* ≤ 0.05, light red and light blue represent the metabolites that increased and decreased respectively at 0.05 ≤ *p* ≤ 0.1, and size of circles represent the magnitude of change. ANOVA contrasts were used to identify metabolites that differed significantly between experimental groups and proper controls, and q-value is reported in Dataset S1.

**Table S1. ANOVA of RT-qPCR and lifespan of *Ubi* > *bmm*-*RNAi* flies.**

**Table S2. ANOVAs of locomotor activity of *Ubi* > *bmm*-*RNAi* flies.**

**Table S3. ANOVAs of sleep of *Ubi* > *bmm*-*RNAi* flies.**

**Table S4. ANOVAs of locomotor activity of *bmm*-*RNAi* lines expressed in fat-body (*Lsp2-Gal4*) and oenocytes (*Dsat1-Gal4* and *OK72-Gal4*) in normal feeding.**

**Table S5. ANOVAs of sleep of *bmm*-*RNAi* lines expressed in fat-body (*Lsp2-Gal4*) and oenocytes (*Dsat1-Gal4* and *OK72-Gal4*) in normal feeding.**

**Dataset S1 (XLSX 598 KB). Metabolome of *Ubi* > *bmm-RNAi* lines.**

**Dataset S2 (XLSX 49 KB). Over-representation analysis of global metabolic pathways for differentially abundant metabolites on *Ubi* > *bmm-RNAi* lines.**

**Dataset S3 (XLSX 175 KB). Overrepresentation analysis of lipids for differentially abundant metabolites on *Ubi > bmm-RNAi* lines.**

**Dataset S4 (XLSX 626 KB). RT-qPCRs and internal control.**

**Dataset S5 (XLSX 37 KB). Raw lifespan data of *Ubi > bmm-RNAi* flies.**

**Dataset S6 (XLSX 86 KB). Average locomotor activity profiles.**

**Dataset S7 (XLSX 17 KB). Average of 24h locomotor activity.**

**Dataset S8 (XLSX 18 KB). Average of locomotor activity during the daytime and nighttime.**

**Dataset S9 (XLSX 150 KB). Individual sleep during the daytime and nighttime.**

**Dataset S10 (XLSX 38.4 MB). Individual activity/sleep bout raw data.**

## References

1. Afshin A, Forouzanfar MH, Reitsma MB, Sur P, Estep K, Lee A, et al. Health effects of overweight and obesity in 195 countries over 25 years. N. Engl. J. Med. 2017; 377:13–27.

2. Saklayen MG. The global epidemic of the metabolic syndrome. Curr. Hypertens. Rep. 2018; 20: 1–8.

3. Locke AE, Kahali B, Berndt SI, Justice AE, Pers TH, Day FR, et al. Genetic studies of body mass index yield new insights for obesity biology. Nature 2015; 518:197–206.

4. Mattson MP. An evolutionary perspective on why food overconsumption impairs cognition. Trends Cogn. Sci. 2019; 23: 200–212.

5. MondaV, La Marra M, Perrella R, Caviglia G, Iavarone A, Chieffi S, et al. Obesity and brain illness: from cognitive and psychological evidences to obesity paradox. Diabetes, Metab. Syndr. Obes. Targets Ther. 2017;10: 473–479.

6. Ugur B, Chen K, Bellen HJ. Drosophila tools and assays for the study of human diseases. Dis. Model. Mech. 2016; 9:235–244.

7. Mattila J, HietakangasV. Regulation of carbohydrate energy metabolism in *Drosophila melanogaster*. Genetics 2017; 207:1231–1253.

8. Heier C, Kühnlein RP. Triacylglycerol metabolism in *Drosophila melanogaster*. Genetics 2018; 210:1163–1184.

9. Musselman LP, Kühnlein RP. Drosophila as a model to study obesity and metabolic disease. J. Exp. Biol. 2018; 221: jeb163881.

10. Musselman LP, Fink JL, Narzinski K, Ramachandran PV, Hathiramani SS, Cagan RL, et al. A high-sugar diet produces obesity and insulin resistance in wild-type Drosophila. Dis. Model. Mech. 2011; 4: 842–849.

11. Reed LK, Lee K, Zhang Z, Rashid L, Poe A, Hsieh B, et al. Systems genomics of metabolic phenotypes in wild-type *Drosophila melanogaster*. Genetics 2014; 197: 781–793.

12. Williams S, Dew-Budd K, Davis K, Anderson J, Bishop R, Freeman K, et al. Metabolomic and gene expression profiles exhibit modular genetic and dietary structure linking metabolic syndrome phenotypes in Drosophila. G3 2015; 5: 2817–2829.

13. Cázarez-García D, Ramírez Loustalot-Laclette M, Markow TA, Winkler, R. Lipidomic profiles of *Drosophila melanogaster* and cactophilic fly species: models of human metabolic diseases. Integr. Biol. 2017; 9: 885–891.

14. Nazario-Yepiz NO, Loustalot-Laclette MR, Carpinteyro-Ponce J, Abreu-Goodger C, Markow TA. Transcriptional responses of ecologically diverse Drosophila species to larval diets differing in relative sugar and protein ratios. PLoS One 2017; 12: e0183007.

15. Zhou S, Morgante F, Geisz MS, Ma J, Anholt RRH, Mackay TFC. Systems genetics of the Drosophila metabolome. Genome Res. 2020; 30: 392–405.

16. Rajan A, Perrimon N. Of flies and men: insights on organismal metabolism from fruit flies. BMC Biol. 2013; 11: 1–8.

17. Palm W, Sampaio JL, Brankatschk M, Carvalho M, Mahmoud A, Shevchenko A, et al. Lipoproteins in *Drosophila melanogaster*—Assembly, function, and influence on tissue lipid composition. PLoS Genet. 2012; 8: e1002828.

18. Gutierrez E, Wiggins D, Fielding B, Gould AP. Specialized hepatocyte-like cells regulate Drosophila lipid metabolism. Nature 2007; 445: 275–280.

19. Grönke S, Müller G, Hirsch J, Fellert S, Andreou A, Haase T, et al. Dual lipolytic control of body fat storage and mobilization in Drosophila. PLoS Biol. 2007; 5: 1248–1256.

20. Bharucha KN, Tarr P, Zipursky SL. A glucagon-like endocrine pathway in Drosophila modulates both lipid and carbohydrate homeostasis. J. Exp. Biol. 2008; 211: 3103–3110.

21. Patel RT, Soulages JL, Hariharasundaram B, Arrese EL. Activation of the lipid droplet controls the rate of lipolysis of triglycerides in the insect fat body. J. Biol. Chem. 2005; 280: 22624–22631.

22. Molaei M, Vandehoef C, Karpac J. NF-κB shapes metabolic adaptation by attenuating Foxo-mediated lipolysis in Drosophila. Dev. Cell 2019; 49: 802–810.

23. Grönke S, Mildner A, Fellert S, Tennagels N, Petry S, Müller G, et al. Brummer lipase is an evolutionary conserved fat storage regulator in Drosophila. Cell Metab. 2005; 1: 323–330.

24. Fischer J, Lefèvre C, Morava E, Mussini JM, Laforêt P, Negre-Salvayre A, et al. The gene encoding adipose triglyceride lipase (*PNPLA2*) is mutated in neutral lipid storage disease with myopathy. Nat. Genet. 2007; 39: 28–30.

25. Wat LW, Chao C, Bartlett R, Buchanan JL, Millington JW, Chih HJ, et al. A role for triglyceride lipase *brummer* in the regulation of sex differences in Drosophila fat storage and breakdown. PLoS Biol. 2020; 18: 1–53.

26. Liu Z, Huang X. Lipid metabolism in Drosophila: development and disease. Acta Biochim Biophys Sin 2013; 45: 44–50.

27. Bi J, Xiang Y, Chen H, Liu Z, Grönke S, Kühnlein RP, et al. Opposite and redundant roles of the two Drosophila: perilipins in lipid mobilization. J. Cell Sci. 2012; 125: 3568–3577.

28. Carvalho M, Sampaio JL, Palm W, Brankatschk M, Eaton S, Shevchenko A. Effects of diet and development on the Drosophila lipidome. Mol. Syst. Biol. 2012; 8: 1–17.

29. Oxenkrug G, Navrotskaya V, Vorobyova L, Summergrad P. Extension of life span of *Drosophila melanogaster* by the inhibitors of tryptophan-kynurenine metabolism. Fly (Austin) 2011; 5: 307–309.

30. Mangge H, Summers KL, Meinitzer A, Zelzer S, Almer G, Prassl R, et al. Obesity-related dysregulation of the Tryptophan-Kynurenine metabolism: role of age and parameters of the metabolic syndrome. Obesity 2014; 22: 195–201.

31. Harbison ST, Yamamoto AH, Fanara JJ, Norga KK, Mackay TFC. Quantitative trait loci affecting starvation resistance in *Drosophila melanogaster*. Genetics 2004; 166: 1807–1823.

32. Harbison ST, McCoy LJ, Mackay TFC. Genome-wide association study of sleep in *Drosophila melanogaster*. BMC Genomics 2013; 14: 281.

33. Jaime MDLA, Hurtado J, Ramirez Loustalot-Laclette M, Oliver B, Markow T. Exploring effects of sex and diet on *Drosophila melanogaster* head gene expression. J. Genomics 2017; 5: 128–131.

34. Rehman N, Varghese J. Larval nutrition influences adult fat stores and starvation resistance in Drosophila. PLoS One 2021; 16: e0247175.

35. Blumrich A, Vogler G, Dresen S, Diop SB, Jaeger C, Leberer S, et al. Fat-body brummer lipase determines survival and cardiac function during starvation in *Drosophila melanogaster*. iScience 2021; 24:102288.

36. Ogilvie RP, Patel SR. The epidemiology of sleep and obesity. Sleep Heal. 2017; 3: 383–388.

37. Noordam R, Bos MM, Wang H, Winkler TW, Bentley AR, Kilpeläinen TO, et al. Multi-ancestry sleep-by-SNP interaction analysis in 126,926 individuals reveals lipid loci stratified by sleep duration. Nat. Commun. 2019; 10: 5121.

38. Vozoris NT. Insomnia symptoms are not associated with dyslipidemia: a population-based Study. Sleep 2016; 39: 551–558.

39. Thompson SN. Trehalose – The insect “blood” sugar. Adv. In Insect Phys. 2003: 31: 205–285.

40. Sayej WN, Paul R, Knight Iii PR, Guo WA, Mullan B, Ohtake PJ, Davidson BA, et al. Advanced glycation end products induce obesity and hepatosteatosis in CD-1 wild-type mice. Biomed Res. Int. 2016; 2016, 1–12.

41. Agh F, Shidfar F. The effects of dietary advanced glycation end products (AGEs) on liver disorders in dietary interventions in liver disease: foods, nutrients, and dietary supplements (eds. Watson, R. R. & Preedy, V. R.) 213–231 (Elsevier Inc., 2019). doi:10.1016/B978-0-12-814466-4.00018-5

42. Pendleton RG, Rasheed A, Sardina T, Tully T, Hillman R. Effects of tyrosine hydroxylase mutants on locomotor activity in Drosophila: a study in functional genomics. Behav. Genet. 2002; 32: 89–94.

43. Yuan Q, Joiner WJ, Sehgal A. A sleep-promoting role for the Drosophila serotonin receptor 1A. Curr. Biol. 2016; 16: 1051–1062.

44. Majeed ZR, Abdeljaber E, Soveland R, Cornwell K, Bankemper A, et al. Modulatory action by the serotonergic system: behavior and neurophysiology in *Drosophila melanogaster*. Neural Plast. 2016; 2016: 7291438.

45. Luo J, Becnel J, Nichols CD, Nässel DR. Insulin-producing cells in the brain of adult Drosophila are regulated by the serotonin 5-HT 1A receptor. Cell. Mol. Life Sci. 2012; 69: 471–484.

46. Nam SB, Kim K, Kim BS, Im H, Lee SH, Kim S, et al. The effect of obesity on the availabilities of dopamine and serotonin transporters. Sci. Rep. 2018; 8: 4924.

47. Wang G-J, Volkow ND, Logan J, Pappas NR, Wong CT, Zhu W, et al. Brain dopamine and obesity. Lancet 2001; 357: 354–357.

48. Dietzl G, Chen D, Schnorrer F, Su K-C, Barinova Y, Fellner M, et al. A genome-wide transgenic RNAi library for conditional gene inactivation in Drosophila. Nature 2007; 448: 151–156.

49. Cichewicz K, Hirsh J. ShinyR-DAM: A program analyzing Drosophila activity, sleep and circadian rhythms. Commun. Biol. 2018; 1: 1–5.

50. Shaw PJ, Cirelli C, Greenspan RJ, Tononi G. Correlates of sleep and waking in *Drosophila melanogaster*. Science 2000; 287:1834–1837.

51. Franz M, Lopes CT, Huck G, Dong Y, Sumer O, Bader GD, Cytoscape.js: A graph theory library for visualisation and analysis. Bioinformatics 2016; 32: 309–311.

52. Bostock M, Ogievetsky V, Heer J. D3: Data-Driven Documents. IEEE Trans. Vis. Comput. Graph. 2011; 17: 2301–2309.

